# A population-based atlas of the human pyramidal tract in 410 healthy participants

**DOI:** 10.1101/251108

**Authors:** Quentin Chenot, Nathalie Tzourio-mazoyer, François Rheault, Maxime Descoteaux, Fabrice Crivello, Laure Zago, Emmanuel Mellet, Gaël Jobard, Marc Joliot, Bernard Mazoyer, Laurent Petit

**Affiliations:** Groupe d’Imagerie Neurofonctionnelle, Institut des Maladies Neurodégénératives - UMR 5293, CNRS, CEA University of Bordeaux, Bordeaux, France; Sherbrooke Connectivity Imaging Lab, University of Sherbrooke, Canada

**Keywords:** White matter anatomy, Pyramidal Tract, Corticospinal tract, Corticobulbar tract, Healthy human, Diffusion imaging, Tractography

## Abstract

With the advances in diffusion MRI and tractography, numerous atlases of the human pyramidal tract (PyT) have been proposed but the inherent limitation of tractography to resolve crossing bundles within the *centrum semiovale* have so far prevented the complete description of the most lateral PyT projections. Here, we combined a precise manual positioning of individual subcortical regions of interest along the descending pathway of the PyT with a new bundle-specific tractography algorithm. This later is based on anatomical priors to improve streamlines tracking in crossing areas. We then extracted both left and right PyT in a large cohort of 410 healthy participants and built a population-based atlas of the whole-fanning PyT with a complete description of its most cortico-lateral projections. Clinical applications are envisaged, the whole-fanning PyT atlas being likely a better marker of corticospinal integrity metrics than those currently used within the frame of prediction of post-stroke motor recovery. The present population-based PyT, freely available, provides an interesting tool for clinical applications in order to locate specific PyT damage and its impact to the short and long-term motor recovery after stroke.

## Introduction

One of the challenges in the human brain connectomic field is to improve our knowledge about major long-range white matter (WM) connections and to link them with behavioral measurements, clinical deficits and recovery (Jbabdi and Behrens, 2012; Jbabdi et al., 2015; Maier-Hein et al., 2017). Among them, the pyramidal tract (PyT) is a long descending pathway which is crucial for the performance of voluntary motricity (Nathan and Smith, 1955; Nyberg-Hansen and Rinvik, 1963; Ebeling and Reulen, 1992). The PyT is generally considered to comprise the corticospinal tract (CST), monitoring movements of the limbs and trunk, and the corticobulbar tract (CBT), directing the movement of the face, head and neck (Nieuwenhuys et al., 2008). Note that the “pyramidal” terminology corresponds to the fact that the PyT runs longitudinally within the pyramids of the *medulla oblongata*, and does not mean that the PyT fibers originate in the pyramidal cells of the cortex.

After its first detailed description (Dejerine and Dejerine-Klumpke, 1901), the pathway of the PyT in the internal capsule has been enriched in a series of papers and accurately localized in the posterior part of the posterior limb of the internal capsule (Englander et al., 1975; Ross, 1980; Kretschmann, 1988; Ebeling and Reulen, 1992). Early neuroimaging studies with T1 and T2 MRI sequences evidenced at this exact location an hypersignal corresponding to the highly myelinated PyT fibers (Curnes et al., 1988; Mirowitz et al., 1989; Yagishita et al., 1994). Recent studies have used this hypersignal as a marker to delineate the CST in order to study its asymmetry (Hervé et al., 2009; Hervé et al., 2011). Below the internal capsule, CST and CBT fibers enter the pons by passing through the feet of the cerebral peduncle (Dejerine and Dejerine-Klumpke, 1901). CST fibers continue their descent in the anterior part of the brainstem, then go along the *medulla oblongata* within the pyramid (Nathan & Smith, 1955) before that 90% of them decussate when entering the spinal cord to finally join their target motoneurons at different levels (Nieuwenhuys et al., 2008). Note that the CBT fibers project to the cranial nerve motor nuclei in the brainstem and most of them do not descend as far as the *medulla oblongata*. However, from a functional point of view (both tracts control voluntary movements) the term “pyramidal tract” has been long used as including corticobulbar as well as corticospinal fibers (Nyberg-Hansen and Rinvik, 1963). Strictly speaking, PyT encompasses all fibers from the cortex to the lower part of the brainstem (Ebeling and Reulen, 1992).

With the emergence of diffusion-weighted magnetic resonance imaging (DWI) and tractography, several atlases of the major white matter pathways have been proposed including rather the CST than the PyT ((Zhang et al., 2010; Thiebaut de Schotten et al., 2011; Yendiki et al., 2011; Rojkova et al., 2016); but see (Archer et al., 2018)). The limitation of DWI-tractography to resolve intricate fiber crossing particularly concerns the PyT for which the most lateral fibers cross within the *centrum semiovale*, the corpus callosum (CC) and the long-range association pathways, mainly the superior longitudinal fasciculus (SLF) and the arcuate fasciculus (AF). As a matter of fact, the previous atlases did not reveal a comprehensive fanning in the lateral projections of the CST/CBT along the primary motor cortex (Thiebaut de Schotten et al., 2011; Rojkova et al., 2016; Archer et al., 2018).

Bürgel et al. (2006) created the first stereotaxic CST atlas by using staining method in 10 adult post-mortem brains. Although they limited their selection to CST fibers originating from the precentral gyrus, they described a comprehensive fanning at the cortical level and the passage of the CST in the posterior part of the posterior limb of the internal capsule (Bürgel et al., 2006). By using diffusion-weighted tractography, Thiebaut de Schotten et al. (2011) proposed a CST atlas in 40 healthy participants. They evidenced dorso-medial CST streamlines in the precentral gyrus but no lateral cortical fanning. This lack was explained by the limitation of the diffusion tensor model in resolving crossing fibers bottleneck between the SLF, CC and PyT streamlines in the *centrum semiovale*. Rojkova et al. (2015) used a more advanced model of tractography (spherical deconvolution), and tracked cortical projections along the central sulcus at the individual level. However, they observed that the lateral projections were too seldom to be represented in the probabilistic atlas. More recently, Archer et al. (2017) constructed a template of the sensorimotor area tracts (SMAT) by using probabilistic tractography. Compared to previous tractography studies, these authors changed the methodology and seeded from multiple sensory-motor areas instead of the precentral area only. They found direct cortico-spinal streamlines from these frontal areas and obtained a better fanning of cortical projections. However, ventro-lateral projections were still lacking in the probabilistic atlas they constituted with 100 healthy participants of the Human Connectome Project (Archer et al., 2018).

The inherent limitation of diffusion-weighted tractography to resolve crossing bundles within the *centrum semiovale* has so far prevented the complete description of the most lateral PyT cortical projections. The aim of the present study is to tackle these limitations to establish a population-based atlas of the whole-fanning PyT by applying a new bundle-specific tractography algorithm based on anatomical priors (Rheault et al., 2017) to improve streamlines tracking in crossing areas. We also defined dedicated anatomical regions of interest (ROIs) at the individual level in order to obtain a specific and sensitive extraction of the PyT in 410 participants of the BIL&GIN database (Mazoyer et al., 2016). We therefore built a PyT atlas with a complete dorso-lateral fanning in each hemisphere. Because damages of the PyT by stroke, tumor, multiple sclerosis, or cerebral palsy can impact the human motor system, one of the stakes of clinical research is to better evaluate initial PyT damages to better predict short and long-term motor recovery (Groisser et al., 2014; Bigourdan et al., 2016). The present PyT atlas may then be a suitable tool in clinical research to investigate these questions.

## Material and methods

### Participants

The present study used diffusion-weighted and T1-weighted images of 410 healthy participants from the BIL&GIN database, balanced in both sex (53% women) and handedness (49% left-handers) with a mean age of 29.9 (± 7.9) years and 15.1 (± 2.6) years of education (Mazoyer et al., 2016). All participants gave written consent prior to participation in the study, which was approved by the local ethics committee (CCPRB Basse-Normandie).

### MRI acquisitions and processing

#### Structural MRI

High-resolution 3D T1 weighted images were obtained using a 3D-FFE-TFE sequence (TR, 20 ms; TE, 4.6 ms; flip angle, 10°; inversion time, 800 ms; turbo field echo factor, 65; Sense factor, 2; field of view, 256 x 256 x 180 mm; isotropic voxel, 1x1x1 mm^3^). For each participant, the line between anterior (AC) and posterior (PC) commissures was identified on a mid-sagittal section, and the T1-MRI volume was acquired after orienting the brain in the bi-commissural coordinate system. Each participant T1-MRI volume was warped to its native diffusion space using ANTS linear and nonlinear registration (Avants et al., 2011).

#### Diffusion MRI

Diffusion-weighted imaging (DWI) data were acquired using a single-shot spin-echo echo-planar sequence composed of one b_0_ map (b = 0 s/mm^2^) followed by 21 diffusion gradient directions maps (b = 1000 s/mm^2^) homogenously spread over a half-sphere and the 21 opposite directions spread over the opposite half-sphere. A second series of one b_0_ and 42 DWI volumes was acquired to increase signal to noise ratio. Seventy axial slices parallel to the AC-PC plane were acquired from the bottom of the cerebellum to the vertex with the following imaging parameters: TR = 8500 ms; TE = 81 ms; angle = 90°; SENSE reduction factor = 2.5; field of view 224 mm; acquisition matrix 112 x 112, 2 x 2 x 2 mm^3^ isotropic voxel. 70 axial slices parallel to the AC-PC plane were acquired from the bottom of the cerebellum to the vertex.

Individual raw DWI data were first corrected for eddy current distortion using the FMRIB software library (Smith et al., 2004) and then processed using Dipy-related tools (Garyfallidis et al., 2014) to obtain the diffusion tensor, the fractional anisotropy (FA), 3-direction vector colored FA (RGB) and fibers orientation distribution of spherical harmonics order 6 (fODF, (Descoteaux et al., 2009)) maps. Note that these different maps were upsampled to 1 mm^3^ spatial resolution using trilinear interpolation (Girard et al., 2014). B_0_, FA and RGB maps were used for the positioning of the regions of interest which delineate the PyT.

### Definition and positioning of ROIs (Figure 1)

In order to extract the PyT of each participant (n = 410), we manually defined and positioned, in each hemisphere, 3 ROIs along its pathway at the level of the internal capsule, the midbrain and the *medulla oblongata*. The manual delineation of these ROIs, their visualization and the filtering of the PyT were performed with TrackVis (http://www.trackvis.org/). Additional anatomical ROIs (referred as JHU-…) were used from the JHU template (Zhang et al., 2010) once warped to the native diffusion space of each subject using ANTS linear and non-linear registration.

**Figure 1.**
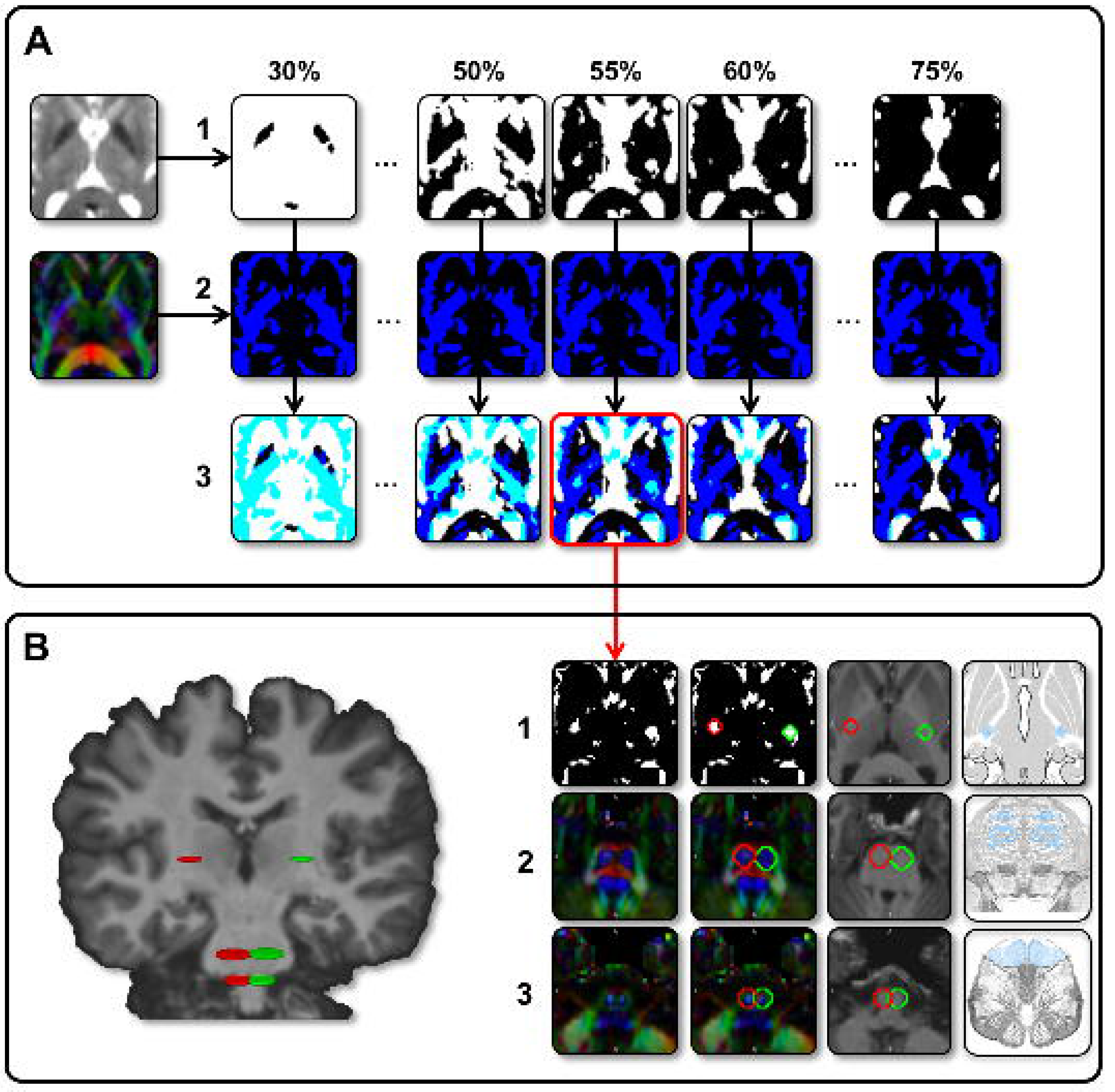
IC-ROI, MB-ROI and MO-ROI definitions. The binarized b_0_-map thresholded from 30 to 75% of maximum value (A1) was multiplied by the binarized RGB-based blue map thresholded at 50% (A2) to build enhanced thresholded-b_0_ maps to highlight the PyT hypersignal at the level of IC (55% in A3-B1). MB was positioned on the blue signal in the anterior part of the midbrain (B2) as opposed to the posterior blue signal corresponding to the lemniscus tract. MO was positioned to contain the pyramids, which is apparent in blue on the RGB map at the level of the chosen axial T1 slice (B3).

#### Internal Capsule ROI (IC-ROI)

The IC-ROI was a disk of 8 mm of diameter (57 voxels) delineated on a single axial slice. To position this ROI in the z-axis, the barycenter of the JHU-thalamus was first calculated and its z-coordinates used because it nearly corresponds to the middle of the adjacent internal capsule in the z-axis. The IC-ROI was then centered for each participant on the hypersignal localized on an enhanced thresholded-b_0_ map (Figure 1). To do so, we first binarized the b_0_ map thresholded between 30 and 75% of its highest value by step of 5%, highlighting the hypersignal at the level of the internal capsule (Figure 1A.1). We also extracted the blue map from the RGB color map, *i.e.* the voxels with a superior-to-inferior main diffusion orientation, thresholded it at 50% of its highest value and binarized it (Figure 1A.2). A series of enhanced thresholded-b_0_ maps were obtained by overlapping these two maps (Figure 1A.3), highlighting the hypersignal as a single cluster at the level of the internal capsule for a given percentage (for example 55% in the example illustrated in Figure 1A) on which the IC-ROI was centered Figure 1B.1). The final positioning of the IC-ROI was visually controlled by superimposition on the T1 map (Figure 1B.1).

#### Midbrain ROI (MB-ROI)

MB-ROI was a disk of 13 mm of diameter (129 voxels) delineated on a single axial slice. It was used to discriminate the PyT (descending motor pathway) localized in the anterior part of the midbrain from the lemniscus tract (ascending sensory pathway) localized in the posterior part of the midbrain. To position this ROI in the z-axis, the RGB color map was used to highlight both tracts in blue color (Figure 1B.2). The axial section in which the two tracts were the most distant from each other was then selected, and the ROI was centered on the blue signal located in the anterior part of the midbrain.

#### Pyramids of the Medulla Oblongata (MO-ROI)

MO-ROI was a disk of 11 mm of diameter (97 voxels) delineated on a single axial slice, which was used to select fibers entering in the *medulla* while not passing through the cerebellum. Note that at this level, the fibers are forming a pyramidal section. To position the MO-ROI in the z-axis, the T1 and RGB color map were used at the level of the *medulla oblongata*. The criterion of selection was that the white matter of the *medulla oblongata* had to be separated from the white matter of the cerebellum and had to form a pyramidal section in the axial T1 map (Figure 1B.3). For each participant, we then positioned the ROI to encompass the pyramids, which appeared in blue on the RGB map at the level of the chosen axial T1 slice (Figure 1B.3).

#### Anatomical location of ROIS in the MNI-space

The three ROIs along the PyT pathway for both hemispheres were warped in the MNI-space by using ANTS linear and non-linear registration. The center of mass of each ROI was calculated for the three axes (x, y, z), for each hemisphere and for each of the 410 participants.

#### Intra-operator reliability for ROI’s positioning

To ensure that the ROIs positioning was robust, we performed a reproducibility analysis in a random sub-sample of 40 out of the 410 participants. ROIs were positioned twice by the same operator after shuffling the participants’ images and their side, using the same methodology. ROIs coordinates were calculated in order to perform an intraclass correlation between the first and the second positioning and to evaluate the reproducibility. The Euclidean distance was also calculated for both positioning.

### Building a PyT template to be used as bundle of interest (Figure 2)

We computed the whole brain tractogram of 39 participants. Following a procedure that has been previously fully described (Hau et al., 2017), each tractogram was built using constrained-spherical deconvolution and particle-filter tractography with anatomical priors (Girard et al., 2014). In brief, each whole tractogram was composed of about 1.5 millions of streamlines seeded from the white/gray-matter interface (10 seeds/voxel, Figure 2A.1) and terminating within the gray matter with a streamline’s length between 10 and 250 mm. We then extracted the PyT in each hemisphere of these 39 participants by using their manually-defined IC-, MB- and MO-ROIs as points of passages and a JHU-fronto-parietal ROI as termination for the streamlines (Figure 2A.2). This later consisted in a single fronto-parietal ROI in each hemisphere defined by merging all frontal and parietal regions of the JHU template (including superficial white matter regions which were merged with their corresponding gyral region). An outlier removal algorithm was applied on each PyT to prune anatomical outlier streamlines (Côté et al., 2015), *i.e* streamlines with multiple curves and loops. Finally, we set a termination-enhanced strategy by keeping the streamlines only ending in the gray matter part of the JHU-fronto-parietal ROI (Figure 2A.3).

**Figure 2.**
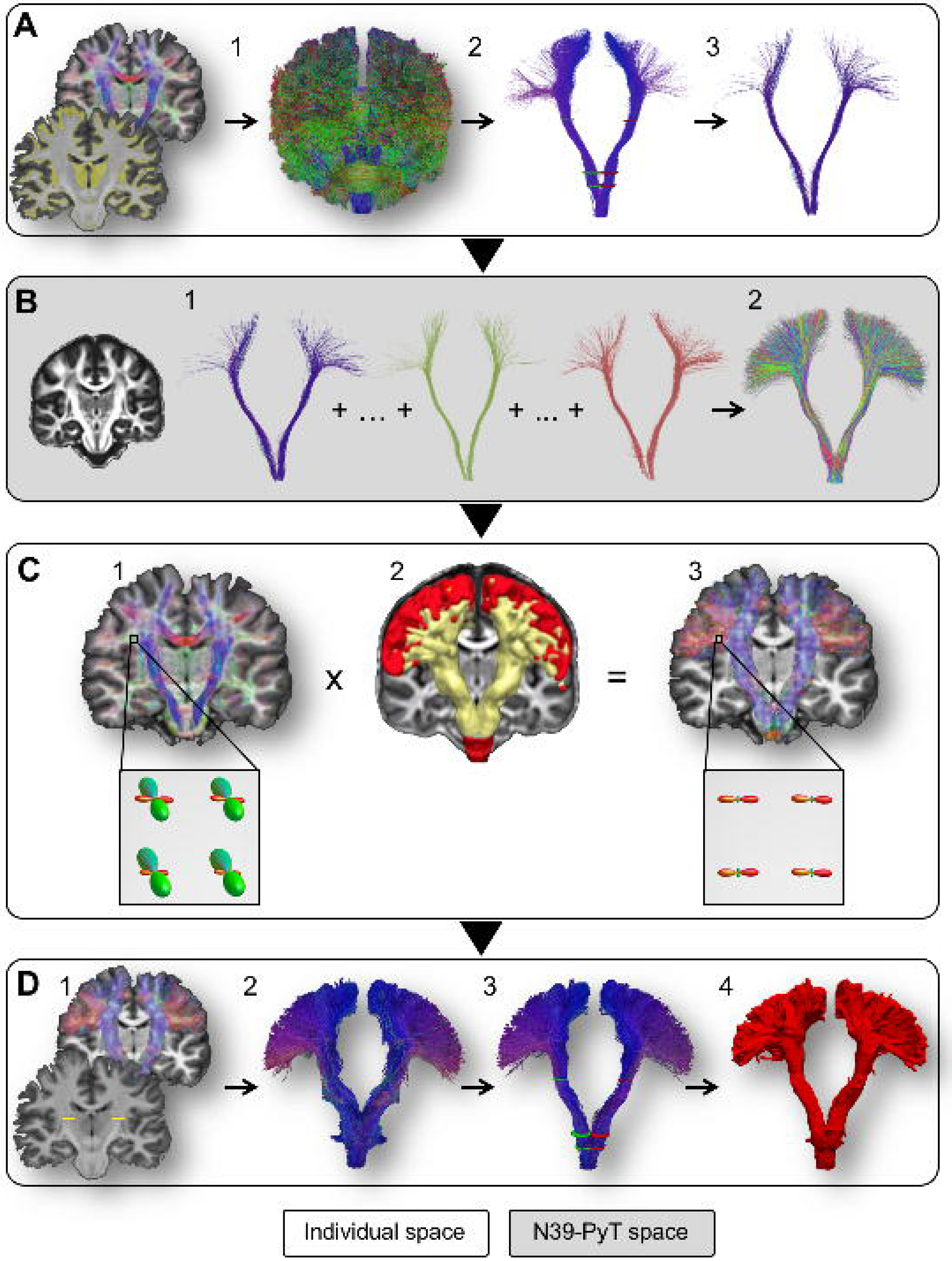
A. Thirty-nine whole-tractograms computed by seeding in the GM/WM interface with a CSD-PFT model (1-2) on which we applied manually defined ROIs (IC-ROI, MB-ROI, MO-ROI) and cleaned the outliers (3). B. N39-PyT template obtained by concatenating the 39 cleaned PyTs in a common space. C. The N39-PyT template (in yellow) used to weight the initial fODF (1) into optimized fODF for PyT tracking within the N39-PyT mask (in red). D. N39-PyT template-specific tractography with the weighted-fODF within the IC-ROI (1-2). Final whole-fanning PyT get through the 3 ROIs manually positioned in 410 participants (3-4).

Second, we build a N39-PyT template from these initial PyTs. We first performed an ANTs template construction (Avants et al., 2011) using the FA maps of the 39 participants to obtain a N39-FA template used as a common space for the N39-PyT template (Figure 2B.1). The nonlinear deformation was then applied to the streamlines of each of the 39 PyTs (Figure 2B.2). Since only the shape and position of the streamlines are relevant for the construction of the PyT template, redundant streamlines were removed using QuickBundles, a hierarchical clustering algorithm and the minimal direct-flip distance operator (Garyfallidis et al., 2012). Each PyT was decimated by discarding non-essential streamlines, *i.e.* those that can be removed with no loss of information about the shape or position of the PyT streamlines (Rheault et al., 2017). This operation helps to reduce in the subsequent optimized tractography the biases caused by an over-representation of straight streamlines in the most medial part of PyT and an under-representation of fanning in the most lateral part of PyT.

### Enhancing orientation distribution within the bundle of interest (Figure 2C)

The N39-PyT template was used as a bundle of interest (BOI) to optimize the final tractography of the PyT in the 410 participants. Since the N39-PyT template was created in a common space, a nonlinear deformation was needed to apply these priors to each participant. We computed the nonlinear deformation matrix between the N39-FA template and the FA map for each of the 410 participants and applied it to the N39-PyT template to displace the N39-PyT streamlines in each individual diffusion space. The next optimization consisted in a voxel-wise weighting of fODF lobes according to the direction of the streamlines from the N39-PyT template (Rheault et al., 2017). The method is based on a track orientation distribution (TOD) map (Dhollander et al., 2014). It produces a new set fODF where the influence of lobes with the same general direction as the PyT is slightly increased, while the influence of lobes with a distinct direction is slightly decreased. This result in a more efficient tracking, with less abrupt changes in direction and a better performance in crossing area resulting in an improved coverage (see Discussion).

### Weighted-fODF tractography of the PyT within the internal capsule ROI (Figure 2D)

At that stage, the streamlines of the N39-PyT template were used as mask to constrain the tracking in the global shape of the PyT (yellow part of Figure 2C-2). A final tractography was then performed by using the particle-filter tractography algorithm within this new tracking mask and the weighted fODF map previously mentioned. The seeding was initiated in the right and left IC-ROI (57 voxels both) with 1000 seeds per voxel. The length of the streamlines was set at a minimum of 80 and a maximum of 180 mm. If not otherwise specified, we used the default parameters. For each of the 410 participants, this created a PyT raw tractogram (Figure 2D-2) on which we applied the 3 manually positioned ROIs to select streamlines (Figure 2D-3). The number of streamlines and the volume in the native space (after binarization, Figure 2D-4) were calculated for each participant.

### PyT atlases

Each individual PyT tractogram was first binarized, then normalized to the MNI space using the ANTS inverse warp and affine matrices that were used to register the JHU template to the diffusion space, to build a population-based atlas of the PyT following the method previously applied to create histology and tractography atlases WM tracts (Bürgel et al., 2006; Thiebaut de Schotten et al., 2011). To do so, the 410 binarized PyT maps were summed and set to a probabilistic map between 0 and 100% overlap (Figure 6).

All the final streamlines of the PyT tractograms of each participant were also normalized to the MNI space in order to concatenate them in a N410-PyT tractogram composed of about 4.4 and 5 millions of streamlines for left and right hemisphere, respectively (Figure 3). A lot of these streamlines once normalized were spatially identical. We thus reduced the number of streamlines composing each N410-PyT by removing redundant streamlines across participants using a hierarchical clustering algorithm and the minimal direct-flip (MDF) distance operator (Garyfallidis et al., 2012).

**Figure 3.**
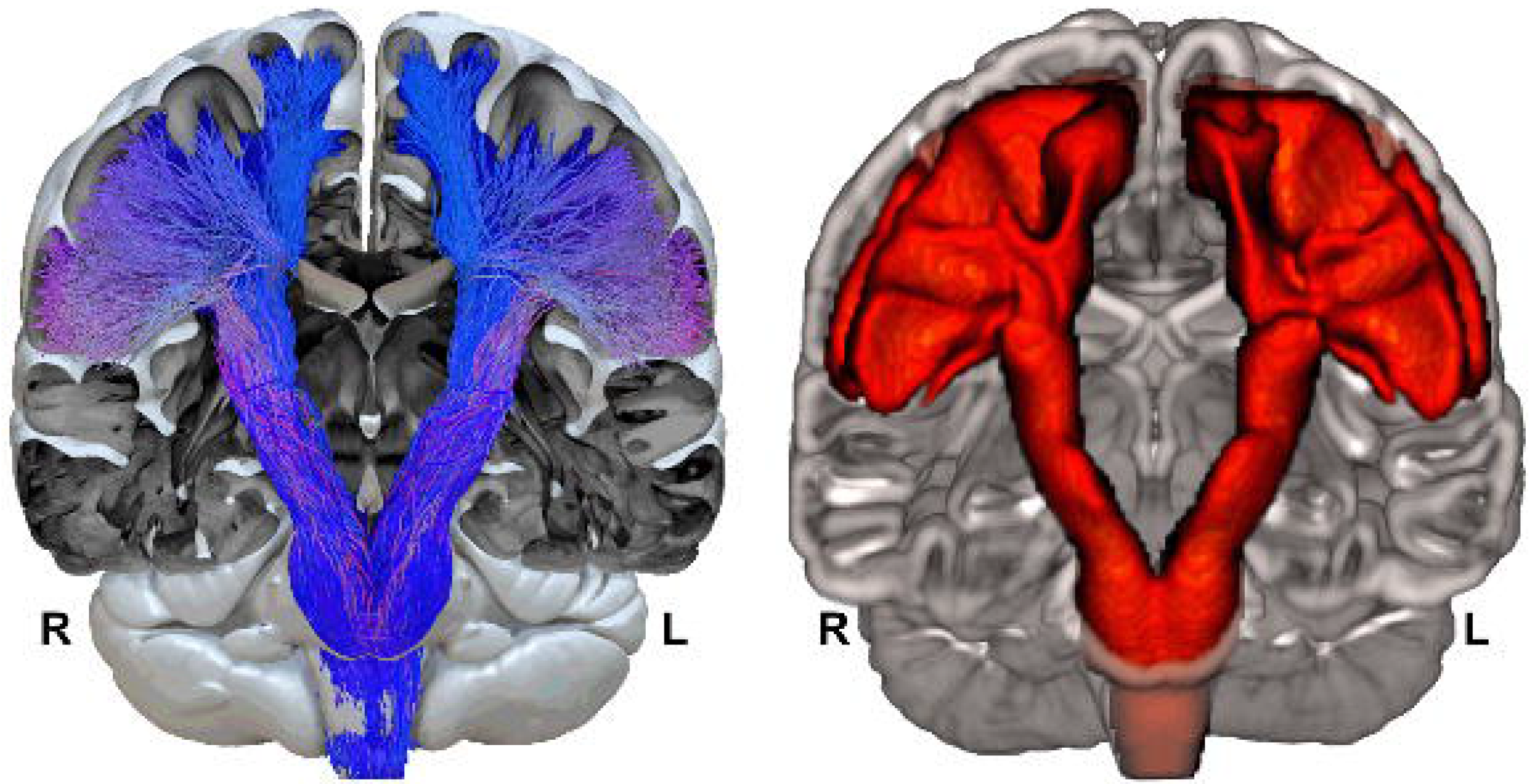
Overview of the whole-fanning PyT atlas (L: left; R: right). On the left side, concatenation of the 410 CSTs warped in the MNI space. Redundant streamlines have been removed using hierarchical clustering algorithm and minimal direct-flip distance operator set at 2 mm. On the right side, tridimensional envelop of the population-based PyT atlas in the MNI space thresholded at 10% (details in Figure 5 and 6). Displays made with Surf Ice (left, www.nitrc.org/projects/surfice/) and MRIcroGL (right, www.nitrc.org/projects/mricrogl/).

### Statistical analysis: PyT asymmetry

Statistical analyses were performed using JMP 11 (www.jmp.com, SAS Institute Inc., 2012). To test the PyT anatomical asymmetry, a repeated MANOVA was computed on 405 participants (note that 5 participants were removed from the analysis because they were inverted left handed). The repeated variable was the hemispheric PyT volume (Left vs Right) in the individual space. Adjustment variables were the Manual Preference (Left *vs*. Right), Gender (Men *vs*. Women), Age and the Cerebral Volume.

## Results

### Mean anatomical localization of the 3 ROIs in the MNI space (Table 1)

Mean coordinates of the center of the IC-ROI were x = -22.3 (standard deviation = 1.1), y = -16.0 (1.3), z = 4.4 (1.1) on the left, and x = 22.8 (1.1), y = -16.0 (1.2), z = 5.0 (1.1) on the right. These coordinates in the MNI space actually correspond to the posterior part of the posterior limb of the internal capsule for both hemispheres.

**Table 1.** Mean (sd) coordinates of the three ROIs along the pathway of the CST in the MNI space (n = 410).

| ROI | MNI coordinates |  |  |
| --- | --- | --- | --- |
|  | x | y | z |
| Left Internal Capsule | -22.32 ± 1.09 | -15.97 ± 1.28 | 4.34 ± 1.13 |
| Right Internal Capsule | 22.81 ± 1.11 | -15.96 ± 1.19 | 4.96 ± 1.11 |
| Left Midbrain | -6.68 ± 1.19 | -22.47 ± 1.85 | -33.86 ± 2.02 |
| Right Midbrain | 7.01 ± 1.13 | -22.60 ± 1.83 | -33.62 ± 2.03 |
| Left Medulla oblongata | -4.59 ± 1.07 | -27.50 ± 1.87 | -43.04 ± 1.89 |
| Right Medulla oblongata | 4.89 ± 1.12 | -27.48 ± 1.79 | -43.21 ± 1.89 |

For the MB-ROI, mean coordinates were x = -6.7 (1.2), y = -22.5 (1.8), z = -33.9 (2.02) on the left and x = 7.0 (1.1), y = -22.6 (1.8), z = -33.6 (2.0) on the right. In the MNI space, these coordinates were actually in the anterior part of the Midbrain.

For the MO-ROI, mean left coordinates were x = -4.6 (1.1), y = -27.5 (1.9), z = -43.0 (1.9) and mean right: x = 4.9 (1.1), y = -27.5 (1.8), z = -43.2 (1.9). In the MNI space, these coordinates correspond to the anterior part of the *medulla oblongata*.

### Intra-operator ROIs positioning reliability (Table 2)

The reproducibility of ROI’s positioning was consistent between the first and second positioning in terms of coordinates. Intraclass correlation coefficients computed were very satisfying being superior to .90 for all ROIs and for all incidences (x, y, z). Mean Euclidean distances between the ROIs were between 0.61 and 1.15 mm, which correspond approximately to a difference of one voxel, which have a 1x1x1 mm^3^ resolution.

**Table 2.** Intraclass correlation coefficients and mean Euclidean distance (standard deviation) between first and second positioning (individual space).

| ROI | X | Y | Z | Euclidean Distance (in mm<sup>3</sup>) |
| --- | --- | --- | --- | --- |
| Left internal capsule | .96 | .99 | .99 | 0.61 ± 0.70 |
| Right internal capsule | .95 | .99 | .99 | 1.02 ± 0.70 |
| Left midbrain | .94 | .99 | .99 | 1.15 ± 0.73 |
| Right midbrain | .92 | .99 | .99 | 1.15 ± 0.77 |
| Left medulla oblongata | .97 | .98 | .99 | 0.99 ± 0.68 |
| Right medulla oblongata | .95 | .99 | .99 | 1.05 ± 0.61 |

### Description of the PyT atlas

Both left and right PyTs were obtained in the 410 participants. Figures 3 shows overall concatenated left and right N410-PyTs. Thanks to the manual positioning of the different ROIs along the PyT pathways in each participant, each of the left and right 410 PyT descended through the *centrum semiovale*, passed through the posterior part of IC, crossed the anterior part of the cerebral peduncle and reached the MO through the anterior part of MB. At the cortical level, we observed a whole-fanning of the PyT around all the convexity of the peri-central cortex. Figure 4 shows the overall distribution of the PyT streamlines in the different frontal and parietal gyri that were initially included in the fronto-parietal JHU ROI (see Methods). The largest proportion of cortical termination was found overwhelmingly in the precentral gyrus (> 50%, Figure 4B), then in the postcentral gyrus (>26%), the superior frontal gyrus (>13%) and to a much lesser extent the superior parietal, supramarginal, middle and inferior frontal gyri. Coronal, axial and sagittal views of these PyT projection per gyri are available in supplementary materials.

**Figure 4.**
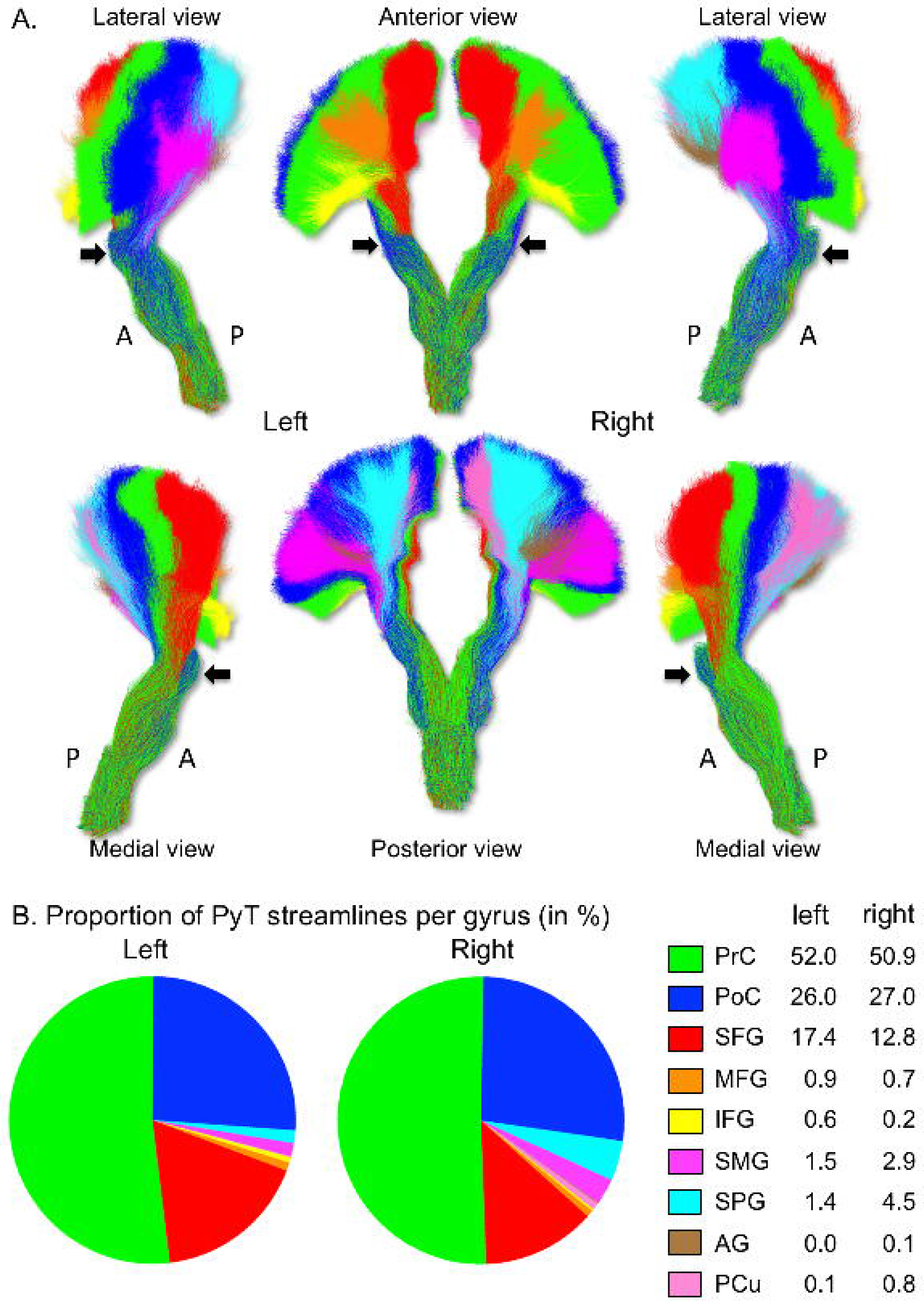
A. Illustration of the distribution of the concatenated PyT streamlines of the 410 participants colored by their gyral termination. B. Proportion of the concatenated frontal and parietal PyT streamlines. Bundle displays made with MI-BRAIN (www.imeka.ca/mi-brain). A: anterior; P: posterior.

It is noteworthy that after a well-ordered cortical organization the streamlines seem to intermingle at the level of the internal capsule while making a twist, the “blueish” streamlines passing in front of the "greenish" streamlines (black arrows in Figure 4A).

#### PyT volume

The analysis of individual hemispheric volumes showed a mean volume of 35775 ± 4002 mm^3^ for the left PyT, and 38055 ± 4380 mm^3^ for the right PyT. The MANOVA revealed that this rightward asymmetry is significant (F=181.0, p < 0.0001). There was no effect of Manual Preference on PyT asymmetry (F = 2.7, p = 0.10), left and right-handers showing the same volume for each hemisphere (Right handed, left hemisphere = 35620 ± 3865; right hemisphere = 37661 ± 4104; Left handed, left hemisphere = 35932 ± 4140; right hemisphere = 38450 ± 4620). As expected, there was an effect of the cerebral volume (F = 75.1, p < 0.0001) with the PyT volume increasing with the cerebral volume. There was an effect of the gender, with men having a bigger PyT volume than women (F = 4.6, p = 0.03; women, left hemisphere = 34333 ± 3803; right hemisphere = 36477 ± 4170; men, left hemisphere = 37274 ± 3645; right hemisphere = 39694 ± 3982) and no age effect (F = 0.1, p = 0.78).

We computed the percentage of overlap between the volume of the PyT extracted before and after the weighting of the fODF maps in each individual. We observe that 91.1 ± 2.0% and 91.6 ± 1.6% of the original left and right PyT volumes overlap with the PyTs built from the weighted-fODF maps. The added volume of PyT fibers obtained with weighted-fODF was in average of 30.8 ± 2.8% and 31.8 ± 2.5% for the left and right hemisphere, respectively. The supplementary figure S1 clearly illustrate that such added volumes correspond to the most lateral fanning parts of the PyTs that were better tracked when applying the present bundle-specific method.

#### Description of the PyT population-based atlas

The PyT atlas had a fanning of cortical projections in the superior, medium and inferior parts of the pre- and post-central gyrus (Figure 5) visible on low (10-30%) as well as high thresholds (70-80%), especially in -12, -16 and -20 coordinates on coronal slices, and in +24 to +60 on axial slices. The superior cortical projections were localized on the posterior coronal planes (-20, -24, -28, -32) whereas the inferior cortical projections were localized more anteriorly (-4, -8, -12, -16). This antero-posterior shift in the dorso-medial direction corresponds to the orientation of the Rolandic sulcus and both precentral and postcentral gyri where the PyT cortical projections were by far found. Figure 6 illustrates the extent of the cortical projection at the different thresholds (from 10% to 90%) and Table 3 lists the volume of the PyT atlas at these different thresholds for both hemispheres. We observed that the right PyT atlas was larger than the left one in the same range whatever the threshold (about 2.7%). Note also that the volumes obtained at the threshold of 40% (left: 35045 mm^3^; right: 37052 mm^3^) were similar to the mean volume observed and the individual level (left: 35775 mm^3^; right: 38055 mm^3^).

**Figure 5.**
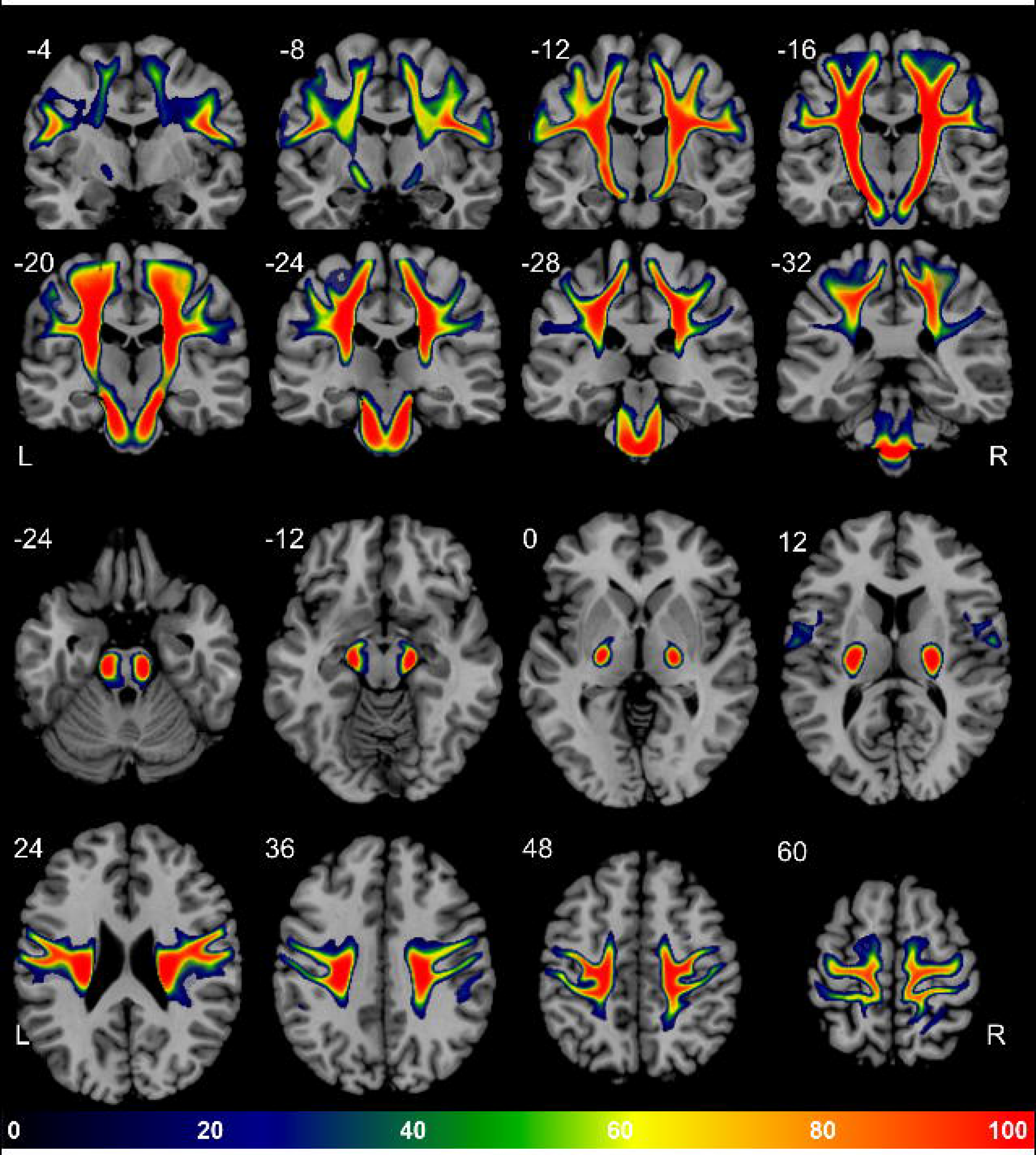
Maps of the PyT atlas in different coronal (top) and axial (bottom) section in MNI space. The color bar indicates the frequency of voxels containing the PyT from 10 to 100% of the 410 participants. For example, a value of 50 in a voxel means that 205 out of 410 individual PyTs are passing through this voxel. L: left; R: right.

**Figure 6.**
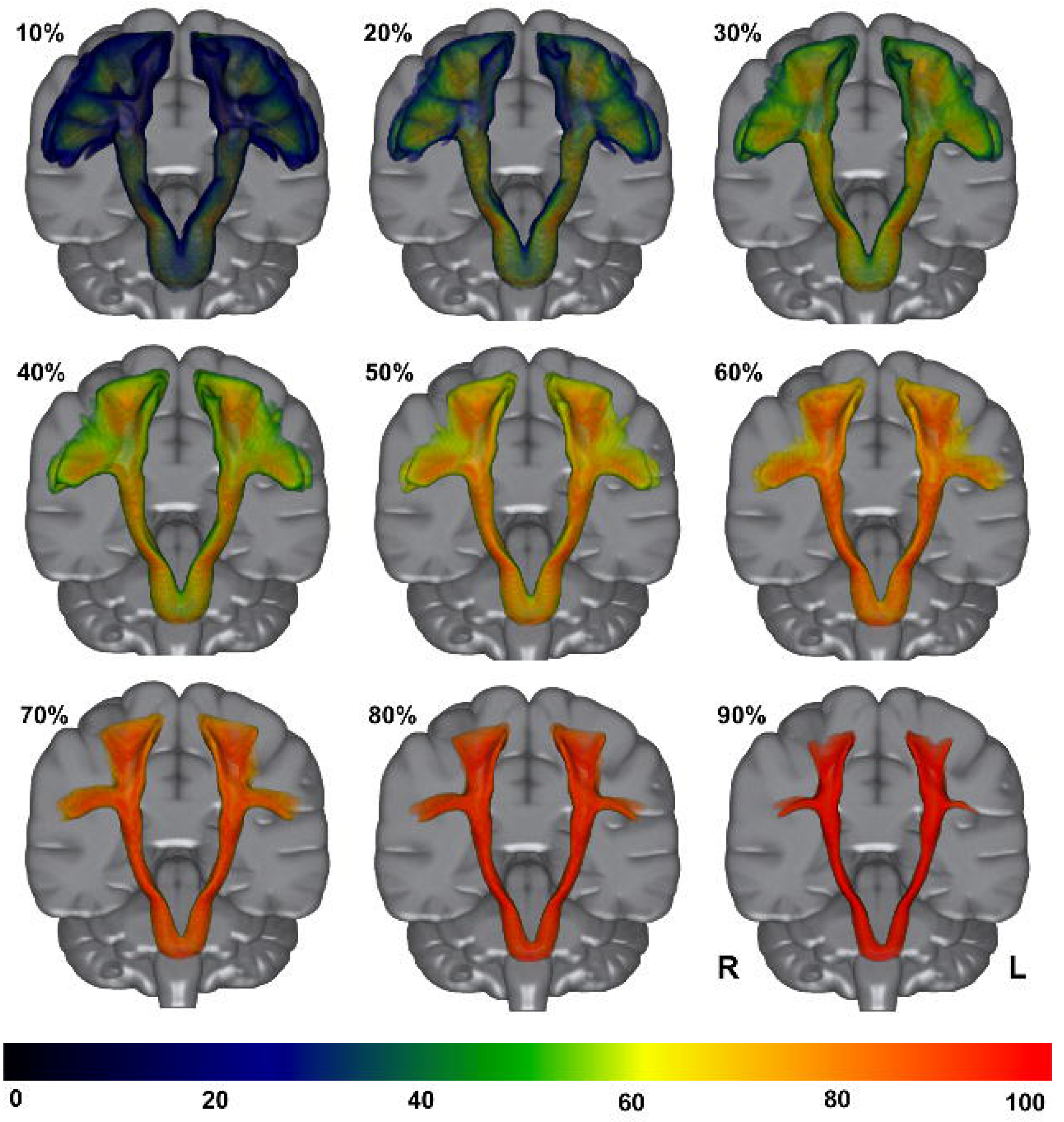
Tri-dimensional frontal view of the population-based PyT volumes at different thresholds from 10% to 90%. The color bar indicates the frequency of voxels containing the PyT from 0 to 100% of the 410 participants. For example, 50% means that 205 out of 410 individual PyTs are included in the volume. L: left; R: right. Displays made with MRIcroGL (www.nitrc.org/projects/mricrogl/).

**Table 3.** Volume of the PyT probabilistic atlas thresholded from 10 to 90% of the 410 participants. For example, 50% means that 205 out of 410 individual PyTs are included in the volume.

| Threshold | Left PyT atlas | Right PyT atlas | (R-L)/(R+L) |
| --- | --- | --- | --- |
|  | (in mm<sup>3</sup>) | (in mm<sup>3</sup>) | (in %) |
| 10 % | 70 706 | 74 271 | 2.5 |
| 20 % | 52 366 | 55 360 | 2.8 |
| 30 % | 42 105 | 44 690 | 2.9 |
| 40 % | 35 045 | 37 052 | 2.8 |
| 50 % | 29 326 | 30 984 | 2.7 |
| 60 % | 24 418 | 25 775 | 2.7 |
| 70 % | 19 912 | 20 971 | 2.6 |
| 80 % | 15 552 | 16 421 | 2.7 |
| 90 % | 10 899 | 11 515 | 2.7 |

As expected, the PyT atlas descends through the *corona radiata* to reach the posterior part of the posterior limb of the internal capsule, between the thalamus and the basal ganglia (axial coordinates 0 and +12, Figure 5). It crosses the cerebral peduncle in its anterior part, the highest thresholds (> 80%) showing that it occupies the central part of the feet of the cerebral peduncle (axial coordinates -12, Figure 5). Lowermost, the PyT atlas is centered in the anterior part of the midbrain until it reaches the *medulla oblongata* (Figure 6).

## Discussion

We first tracked both left and right PyT in a large cohort of 410 healthy participants by combining a precise manual positioning of specific ROI along its descending pathway with a new approach of bundle-specific tractography. Then we built a population-based PyT atlas from this large cohort with a complete description of its most cortico-lateral projections while previous comparable atlases were restricted to its most medio-dorsal part (Catani and Thiebaut de Schotten, 2008; Zhang et al., 2010; Thiebaut de Schotten et al., 2011; Rojkova et al., 2016; Archer et al., 2018).

### PyT cortical projections

We observed a complete fanning following the central sulcus curvature from the medio-dorsal to the ventro-lateral part, closely similar to the one previously described in the dissection and in the histology literature (Dejerine and Dejerine-Klumpke, 1901; Ebeling and Reulen, 1992; Bürgel et al., 2006; Nieuwenhuys et al., 2008) and covering the entire sensori-motor homunculus originally described by Penfield (Penfield and Boldrey, 1937). The presence of a large amount of ventro-lateral projections was striking considering the difficulty of tracking in the *centrum semiovale* reported in the literature (Farquharson et al., 2013), where multiple bundles are crossing. This observation is likely to result from the use of the CSD-PFT algorithm (Girard et al., 2014), but especially with the weighting of the fODF based on a PyT template, which allowed a better tracking of PyT streamlines in the *centrum semiovale* (Rheault et al., 2017). These ventro-lateral projections represent a significant improvement compared to previous CST/PyT extracted using individual tractography (Kumar et al., 2009; Zolal et al., 2012; Groisser et al., 2014; Angstmann et al., 2016; Bigourdan et al., 2016) and also compared to the different previous probabilistic CST/PyT atlases (Zhang et al., 2010; Thiebaut de Schotten et al., 2011; Rojkova et al., 2016; Archer et al., 2018).

We also observed that the PyT cortical projections, namely the corticospinal and corticobulbar tracts, originated in areas beyond the primary motor cortex (M1) along the Rolando sulcus, and even beyond the precentral gyrus. It is consistent with previous studies in non-human primates which showed that the PyT originates from M1 but also from premotor and parietal areas (Armand, 1982; Dum and Strick, 1991; Galea and Darian-Smith, 1994). In humans, the exact extent of PyT cortical origins is still under question (Archer et al., 2018). A post-mortem study has estimated that the majority of PyT fibers (40-60%) originates from the precentral gyrus in human (Jane et al., 1967; Ebeling and Reulen, 1992), a proportion that we also observed since the most represented PyT projection also originated by far from the precentral gyrus in each hemisphere (Figure 4B). The remainder projections first concerned both postcentral and superior frontal gyri then, to a lesser extent, the posterior parts of the middle and inferior frontal gyri, the supramarginal and superior parietal gyri. In their recent study, Archer et al. (2017) addressed this issue in a different manner by tracking descending sensorimotor tracts from motor and premotor ROIs defined by functional brain imaging (Mayka et al., 2006), hinting that both CST and CBT cortical origins certainly extended beyond M1. Similar results were originally described using functionally-defined motor and premotor seedings (Seo and Jang, 2013). The extent of cortical PyT origins is an important question since the existing CST/PyT atlases choose to restrict the cortical origins to the precentral gyrus (Bürgel et al., 2006; Zhang et al., 2010; Thiebaut de Schotten et al., 2011; Rojkova et al., 2016), which does not seem to correspond to the exact anatomical definition of the PyT. Further investigations including blunt Klingler microdissection ideally combined with diffusion-weighted tractography are needed to investigate this issue.

### PyT anatomical course

As expected since the seeding IC-ROI was positioned at that location in each of the 410 participants, the present PyT atlas crossed the posterior part of the posterior limb of the internal capsule as evidenced in previous dissection studies (Englander et al., 1975; Ross, 1980; Kretschmann, 1988). It is noteworthy that its mean location both in terms of the mean coordinates of the IC-ROI and the PyT map (Figure 5) was exactly the same that in the microscopically defined stereotaxic atlas of the CST (restricted to the M1 projection) (see Figure 7 in (Bürgel et al., 2006)).

We observed that after a well-ordered cortical organization the streamlines were intermingled at the level of the internal capsule while making a twist, the “blueish” medial streamlines with a more medial origin passing in front of the “greenish” streamlines (black arrows in Figure 4A). This is very consistent with previous anatomical description of the PyT course (Nathan and Smith, 1955). These authors wrote *“From the precentral gyrus, the fibers come together to form a band on the internal capsule. To narrow down to pass through a band-like gap, the fibers undergo a screw-like gyration, the fibers from the inferior part of the gyrus coming to lie most medially and those from the paracentral lobule coming to lie most laterally. (pg. 264)”* The descending course of the PyT atlas into the anterior part of midbrain was also similar to the pathway reported in dissection and histology studies (Dejerine and Dejerine-Klumpke, 1901; Nathan and Smith, 1955; Bürgel et al., 2006). Note that thanks to the accurate ROI positioning in the midbrain, the PyT atlas correctly crossed the feet of the cerebral peduncle which constituted a validation of the PyT pathway, since no ROI was defined at this level to restrict the fibers’ pathway.

Our PyT atlas showed a higher resolution in its sub-cortical pathway than previous CST/PyT atlases (Thiebaut de Schotten et al., 2011; Rojkova et al., 2016; Archer et al., 2018). This higher accuracy was due to the manual positioning of 3 sub-cortical ROIs in each of the 410 participants (posterior part of the posterior limb of the internal capsule; anterior part of the midbrain and pyramids in the *medulla oblongata*), whereas previous studies applied automated positioning tools and a lower number of sub-cortical ROIs.

### PyT asymmetry

To our knowledge, this is the first report of a rightward PyT volume asymmetry. Previous tractography studies that have quantified the CST/PyT volume in healthy individuals found either an absence of asymmetry or a leftward asymmetry. Kwon et al. (2011) extracted the CST involved in upper limb motricity (hands and fingers) and inferior limb motricity (legs and feet) in 43 right-handed healthy adults using a probabilistic tractography algorithm. Their results showed an absence of asymmetry (Kwon et al., 2011). Kumar et al. (2009) computed a complete tractogram using a deterministic algorithm in 42 children (3-17 years, 33 right-handed). They traced a first ROI in the midbrain and a second in the *corona radiata* and reported a significant leftward asymmetry of the volume of the CST, with no effects of manual preference, and, contrary to the present result, no effect of sex (Kumar et al., 2009). However, the use of a deterministic algorithm limited the cortical projections of their CST in the area corresponding to lower limbs motricity. By using a probabilistic tractography algorithm, Seo & Jang (2011) found no asymmetry in 36 healthy right-handed adults, except for a sub-component originating from the supplementary motor area that was leftward asymmetrical (Seo and Jang, 2013). The discrepancy between these previous studies may be due to methodological limitations such as differences in terms of ROIs choice and reconstructed CST-PyT with few or no ventro-lateral cortical projections that would impact on the estimation of the tract volume. The rightward asymmetry we presently report has the advantage of coming from a much larger sample of participants.

Finally, the right-ward PyT asymmetry cannot be explained by handedness. The large and balanced number of right and left-handers in the BIL&GIN database is what makes this negative result important for the community since it cannot be attributed to a lack of statistical power. Interestingly, a recent study based on HCP data (557 right-handers, 49 left-handers) showed that the handedness cannot explain the different asymmetries observed for anisotropy measures between left and right CST (Volz et al., 2018). Of note, PyT atlases of right- and left-handers have been also released for future studies that would benefit from higher specificity (http://www.gin.cnrs.fr/en/tools/).

### Limitations

One may first consider that the number of directions and b-value of the present DWI data would be far from optimal for the original ODF estimation. The BIL&GIN data have been acquired in 2010-2011 while the standard number of gradient direction maps in diffusion MRI was not intended to perform current up-to-date tractography. But we originally optimized the acquisition protocol in order to improve the signal to noise ratio (SNr). Actually, the DWI data were acquired with 21 directions homogenously spread over a half-sphere, then doubled with the acquisition of 21 opposite directions spread over the opposite half-sphere, and these two series of acquisition were again duplicated, meaning that we acquired four times the 21 gradient direction maps that were therefore averaged improving the SNr by a factor 2. The quality of the present DWI data and the excellent SNr allowed us to instantiate the fODF reconstruction of high-resolution spherical harmonics order 6 with constrained-spherical deconvolution (Descoteaux et al., 2009), allowing a good estimation of 28 coefficients even from 21 measurements resulting in a good angular contrast (see Supplementary Materials for detail about the quality control of the fODF maps). Thank to these optimizations for the acquisition protocol and modeling of the diffusion metrics, we were currently able to apply cutting-edge tractography algorithm on such “old” data. Our previous publications combining fODF-based tractography of the same type of dMRI data and microdissection validation have documented such an improvement and validate our methodological approach since we were able to revisit the anatomy of the uncinate fasciculus (Hau et al. 2017) and to describe the homotopic and heterotopic connectivity of the frontal part of the corpus callosum (De Benedictis et al. 2016).

Although the manual ROIs placement allows a fine-grained and specific extraction of the PyT, one may notice that we used ROIs with the same size in all participants. While this was the only way to complete a rigorous seeding in the internal capsule, it could be considered as a limitation considering individual variability in brain size. We also have to consider the inherent limits in spatial resolution of tractography: the present tractograms were at a millimeter scale (1x1x1 mm^3^) while axons are at a micro-millimeter scale, meaning that one voxel can contain millions of fibers. As a result, quantification in tractography (whether it is the number of streamlines or the volume of a bundle) has to be supported by anatomical data such as post-mortem dissection or histology studies to be validated (Jones et al., 2013; Maier-Hein et al., 2017). The fact that the descending subcortical pathway of the present PyT atlas is highly similar to histological atlas from Bürgel et al. (2006) can be considered as a first validation step. However, Bürgel’s study was restricted to the precentral origin of the CST/PyT that probably underestimated the complete cortical extent of the human PyT.

## Conclusion

In the present study, we built a population-based atlas of the PyT in 410 healthy participants by combining (1) manual positioning of sub-cortical regions of interest and (2) an advanced tractography methodology that allowed to track the whole-fanning PyT. We then elaborated the first atlas of *pyramidal tract* having both sub-cortical accurate anatomical pathway and comprehensive fanning projections along the central sulcus. Further works combining diffusion with functional imaging will permit to identify the sub-components of the PyT related to upper limbs motricity, lower limbs motricity or mouth, tongue and face motricity as typically defined in the *homunculus* (Penfield and Boldrey, 1937). The present population-based PyT atlas provides an interesting tool for clinical applications in order to locate specific PyT damage and its impact to the short and long-term motor recovery after stroke (Groisser et al., 2014; Bigourdan et al., 2016; Archer et al., 2018). The whole population-based PyT atlases (N=410), as well as specific PyT atlases for right- and left-handers, are freely available at http://www.gin.cnrs.fr/en/tools/.

## Supplementary Figures

Supplementary Figure 1. Additional volume of PyT (in blue) obtained in average in the 410 participants when applying the bundle-specific method to extract the PyT. In green, the mean common PyT volume obtained before and after the weighting of the fODF maps. All data overlapped on the coronal view of the MNI-152 anatomical template passing through the population-based PyT atlas. L: left, R: right.

**Supplementary Figure 2**. Coronal, axial and sagittal plates of the concatenated PyT streamlines of the 410 participants (see Figure 3) colored by their frontal and parietal cortical termination.

## Acknowledgments

We are grateful to Dr. Thomas Tourdias for helpful comments and discussion.

